# Development of an imaging toolbox to assess the therapeutic potential and biodistribution of macrophages in a mouse model of multiple organ dysfunction

**DOI:** 10.1101/372482

**Authors:** Jack Sharkey, Lorenzo Ressel, Nathalie Brillant, Bettina Wilm, B. Kevin Park, Patricia Murray

## Abstract

Cell-based regenerative medicine therapies require robust preclinical safety, efficacy, biodistribution and engraftment data prior to clinical testing. To address these challenges, we have developed an imaging toolbox comprising multi-spectral optoacoustic tomography and ultrasonography, which allows the degree of kidney, liver and cardiac injury and the extent of functional recovery to be assessed non-invasively in a mouse model of multi-organ dysfunction. This toolbox allowed us to determine the therapeutic effects of adoptively transferred M2 macrophages. Using bioluminescence imaging, we could then investigate the association between amelioration and biodistribution. Macrophage therapy improved kidney and liver function to a limited extent, but did not ameliorate histological damage. No improvement in cardiac function was observed. Biodistribution analysis showed that macrophages homed and persisted in the injured kidneys and liver, but did not populate the heart. Our data suggest that the limited improvement observed in kidney and liver function could be mediated by M2 macrophages.

## Introduction

Cell-based regenerative medicine therapies (RMTs), which include pluripotent stem cells, mesenchymal stromal cells and macrophages, have the potential to treat a variety of human diseases. However, before these therapies can be routinely used in the clinic, accurate information regarding their safety and efficacy should be obtained from appropriate preclinical models. It is also important to gain an understanding of the therapeutic mechanisms of the RMTs, such as whether their ability to ameliorate injury is dependent on their engraftment in the damaged tissues. Issues which currently prevent the generation of such data include (i) the limitations associated with commonly used blood biomarkers of organ injury, such as serum creatinine (SCr) for renal function (Molitoris et al., 2007, Kellum et al., 2002, Ferguson et al., 2008, Chertow et al., 2005,Bonventre et al., 2010), and alanine aminotransferase for liver function (Vanderlinde, 1986, Ozer et al., 2010,Marrer and Dieterle, 2010); (ii) the technical limitations in repeated blood and urine sampling in small rodents; and (iii) the difficulties associated with monitoring organ function in small animal species longitudinally. The assessment of organ injury in small rodents classically involves measurements of serum or urine biomarkers, or histopathological analysis. Since the latter is usually only undertaken at post-mortem, it fails to allow the progression of injury to be monitored in the same animals longitudinally, requiring animals to be culled at multiple time points, which is not in keeping with NC3Rs’ principles and also reduces the power of the statistical tests. Furthermore, current methods make it difficult to assess the safety, efficacy and therapeutic mechanisms of cell-based RMTs in comorbid conditions where more than one organ is affected, such as in the cardiorenal and hepatorenal syndromes (Mayfield et al., 2016, Gonzalez-Calero et al., 2014,Erly et al., 2015).

In this current study we set out to develop a multimodal imaging strategy to monitor the function of the liver, kidney and heart longitudinally in BALB/c mice in a single imaging session. We utilised multispectral optoacoustic tomography (MSOT) to assess kidney and liver function, and traditional ultrasound (US) measurements to assess cardiac function. MSOT is a technique which uses multiple excitation wavelengths to resolve specific sources of absorption, whether they are endogenous or exogenous (Taruttis et al., 2012). It relies on thermoelastic expansion which occurs when energy from a laser capable of emitting light at a range of wavelengths is absorbed by molecules within the tissues. This causes electrons to move to an excited state, generating heat and a resultant pressure wave which is detected by an ultrasound detector. The specific absorption profile of endogenous or exogenous molecules allows their identification within living animals in a minimally invasive manner (Comenge et al., 2018). We have previously described methods of monitoring kidney or liver function using MSOT in models of chronic kidney injury and acute liver injury (Brillant et al., 2017,Scarfe et al., 2015).

Here, we utilised an acute model of adriamycin (ADR) - (doxorubicin-) induced multi-organ injury and assessed organ function on days one and four post drug administration in order to determine the extent of disease progression. ADR is an anthracycline antibiotic which is clinically administered as a chemotherapeutic agent; however, its use is limited predominantly due to cardiotoxicity mediated through a number of mechanisms. In rodents it can induce both chronic and acute kidney injury, as well as cardiac and hepatic dysfunction (Roomi et al., 2014, Saad et al., 2001,Scarfe et al., 2015).

To test the effectiveness of this MSOT-US bi-modal imaging strategy for monitoring the ameliorative potential of cell-based RMTs, after recording the extent of kidney, liver and cardiac dysfunction on day one, mouse bone marrow-derived alternatively activated (M2) macrophages (BMDMs) were administered intravenously into BALB/c mice and their ability to improve the function of the aforementioned organs was monitored on day four and compared with a saline placebo. M2 BMDMs were used in this study because previous reports have already demonstrated that they can ameliorate kidney injury, liver injury and cardiac injury (Wang et al., 2008, Bai et al., 2017,Shiraishi et al., 2016). However, as far as we are aware, M2 BMDMs have not previously been assessed for their ability to ameliorate the injury of multiple organs simultaneously. We then set out to assess the relationship between the biodistribution of the M2 BMDMs and their ability to ameliorate injury in each of the three organs by utilising bioluminescent imaging of M2 BMDMs expressing luciferase.

## Results

### Use of high frequency ultrasonography to investigate the effect of BMDMs on cardiac function in ADR-dosed mice

High frequency ultrasonography was used to determine whether cardiac function was affected in treated (ADR+BMDM), injured (ADR) and control mice (Fig 1) by measuring the following functional parameters: fractional shortening (FS), ejection fraction (EF), stroke volume (SV) and cardiac output (CO). The mean ΔFS (Fig 2A) and ΔEF (Fig 4B) was decreased in the ADR and ADR+BMDM groups when compared to uninjured controls, but not significantly so. However, there were significant changes in ΔCO (Fig 2C) and ΔSV (Fig 2D), with both parameters being significantly reduced in the ADR and ADR+BMDM groups compared to healthy controls.

**Figure 1:**
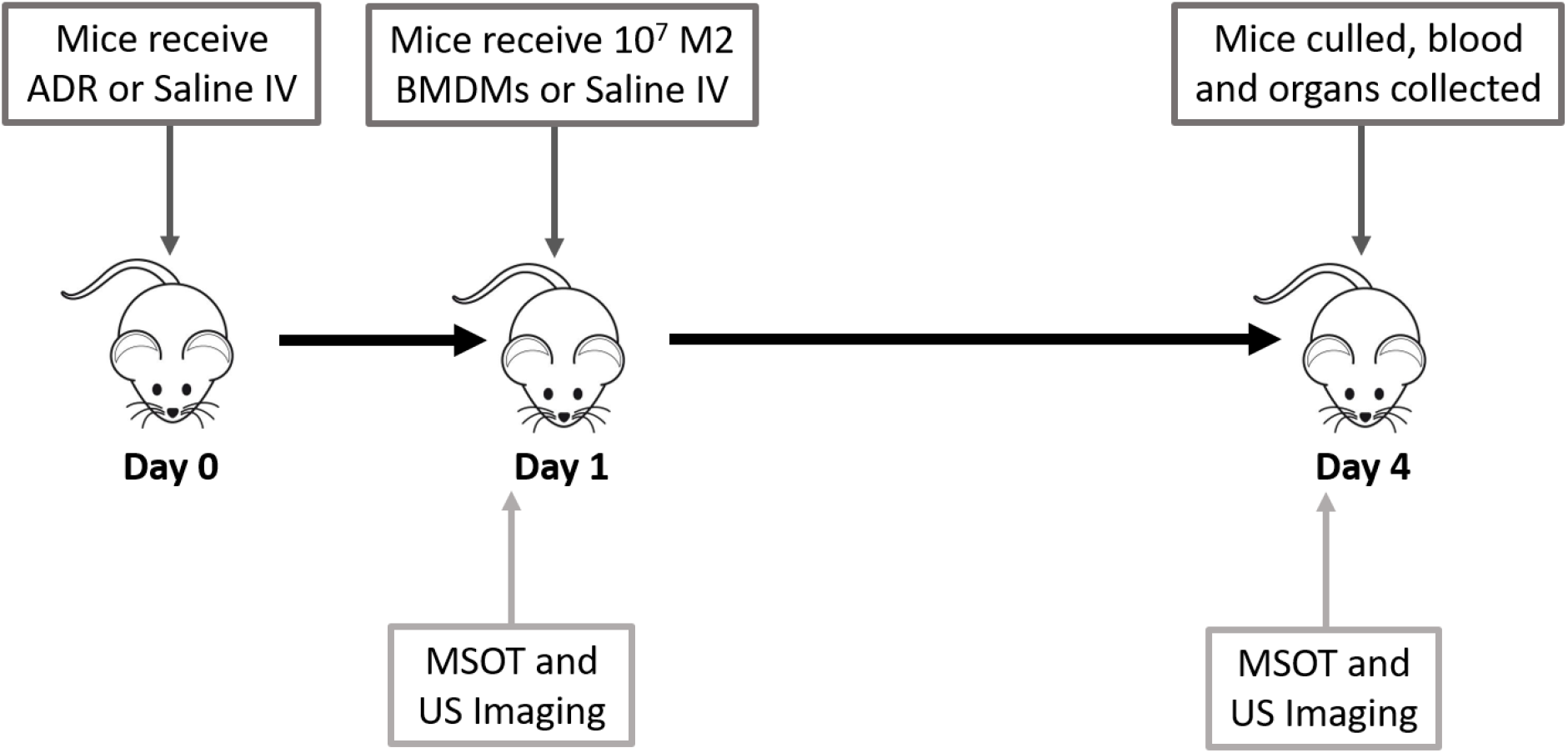
Schematic showing the experimental protocol.

**Figure 2:**
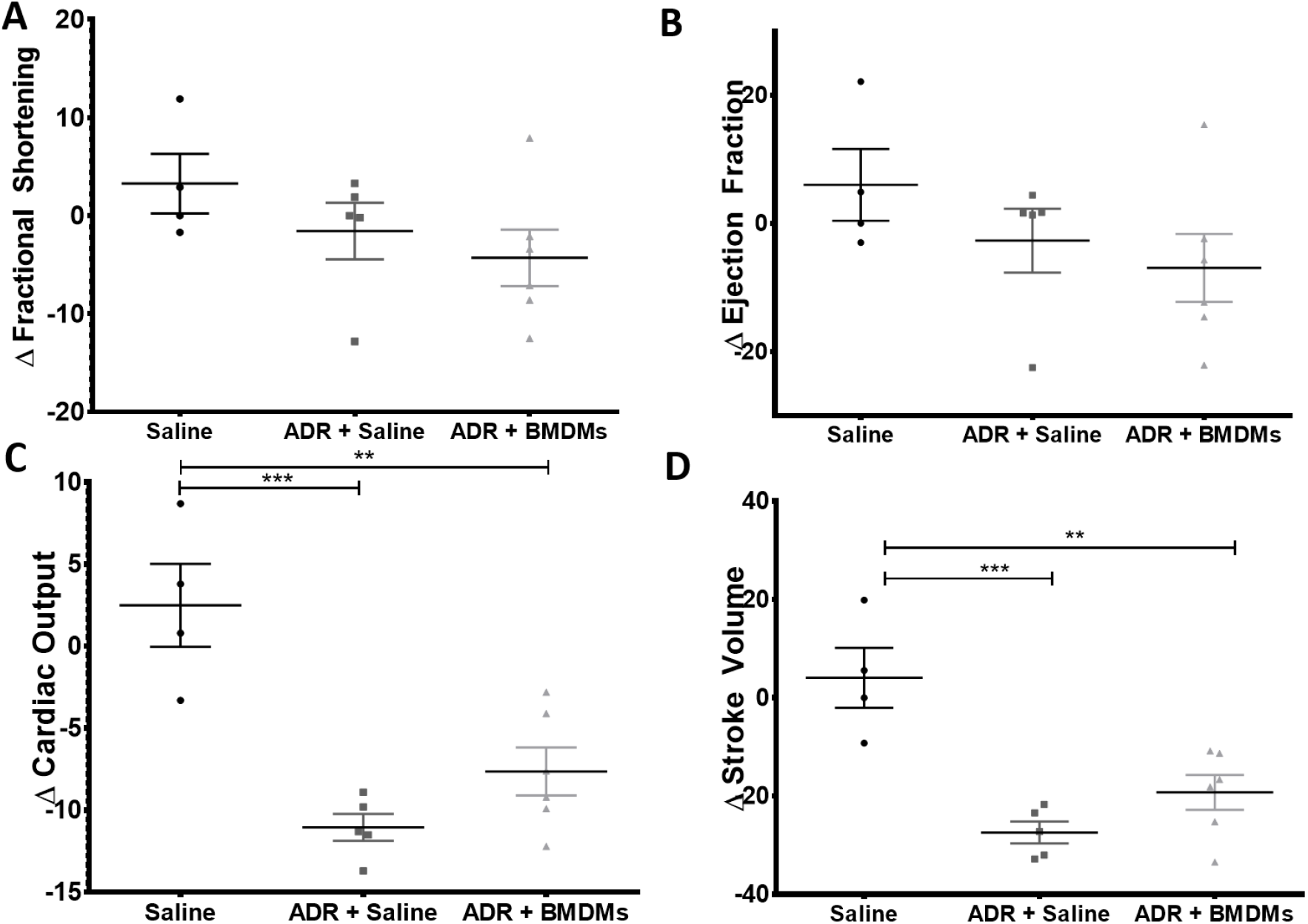
Cardiac parameters as measured by ultrasound. Fractional shortening (A), ejection fraction (B), cardiac output (C) and stroke volume (D) were quantified. Each parameter is represented as the change in the parameter between days 1 and 4 in each mouse. Each data point represents an individual mouse. ** P<0.01, *** P<0.001.

### Use of MSOT to investigate the effect of BMDMs on renal function in ADR-dosed mice

MSOT was used to assess renal function in healthy mice, those which received ADR and those which received ADR+BMDMs, on days 1 and 4 following ADR administration (Fig 1). Uninjured control mice lost approximately 2.5% of their bodyweight over the 4 day time-course of the experiment whereas animals that received ADR lost approximately 18% of their bodyweight, irrespective of whether they were administered macrophages (Sup Fig 2, Table 1). Kidney function was assessed using MSOT, as previously described (Scarfe et al, 2015), to measure the renal clearance of the near infrared dye, IRDye 800 carboxylate (IRDye) which is exclusively filtered by the kidney (Sup Fig 1). Regions of interest were drawn around the cortico-medullary region and the renal pelvis to generate intensity data for quantitative measurements (Fig 3A). We observed that the typical clearance curves of IRDye on day 4 in both cortex and pelvis were changed in animals with ADR-induced kidney injury when compared to those of healthy control mice (Fig. 3B, C). Specifically, instead of a single peak and subsequent decay in the cortex, the curve peaked, dropped but then rose again and slowly decreased over time. In the pelvis, a peak was observed at the same time point as in the cortex, followed by a second peak and subsequent rise in the signal intensity. The clearance kinetics of IRDye in mice after ADR injury and BMDM administration were also altered compared to the healthy controls (Fig 3D). To assess renal function we utilised the ratio of the AUC for the cortex and pelvis (AUC C:P) rather than the Tmax delay (difference in time at which signal was at maximum between cortex and pelvis) or cortex exponential decay time used previously (Scarfe et al, 2015). For quantitative analysis of the changes observed in each individual animal, IRDye clearance data were expressed as the difference in the AUC Cortex:Pelvis between the measurements taken on days 1 and 4 (ΔAUC C:P). There was a significant increase in the ΔAUC C:P between healthy and the ADR group, but not between healthy and the ADR+BMDM group (Fig 3E). In addition, we assessed renal function using classical markers of blood urea nitrogen (BUN) and serum creatinine (SCr) on day 4 (Fig 3F, G respectively). BUN was significantly elevated in the ADR and ADR+BMDM groups compared to healthy controls, whereas there were no significant changes in SCr between all three treatment groups. Furthermore, there was a significant correlation between the AUC C:P and day 4 BUN measurements (Fig 3H, P=0.013, R^2^=0.39).

**Figure 3:**
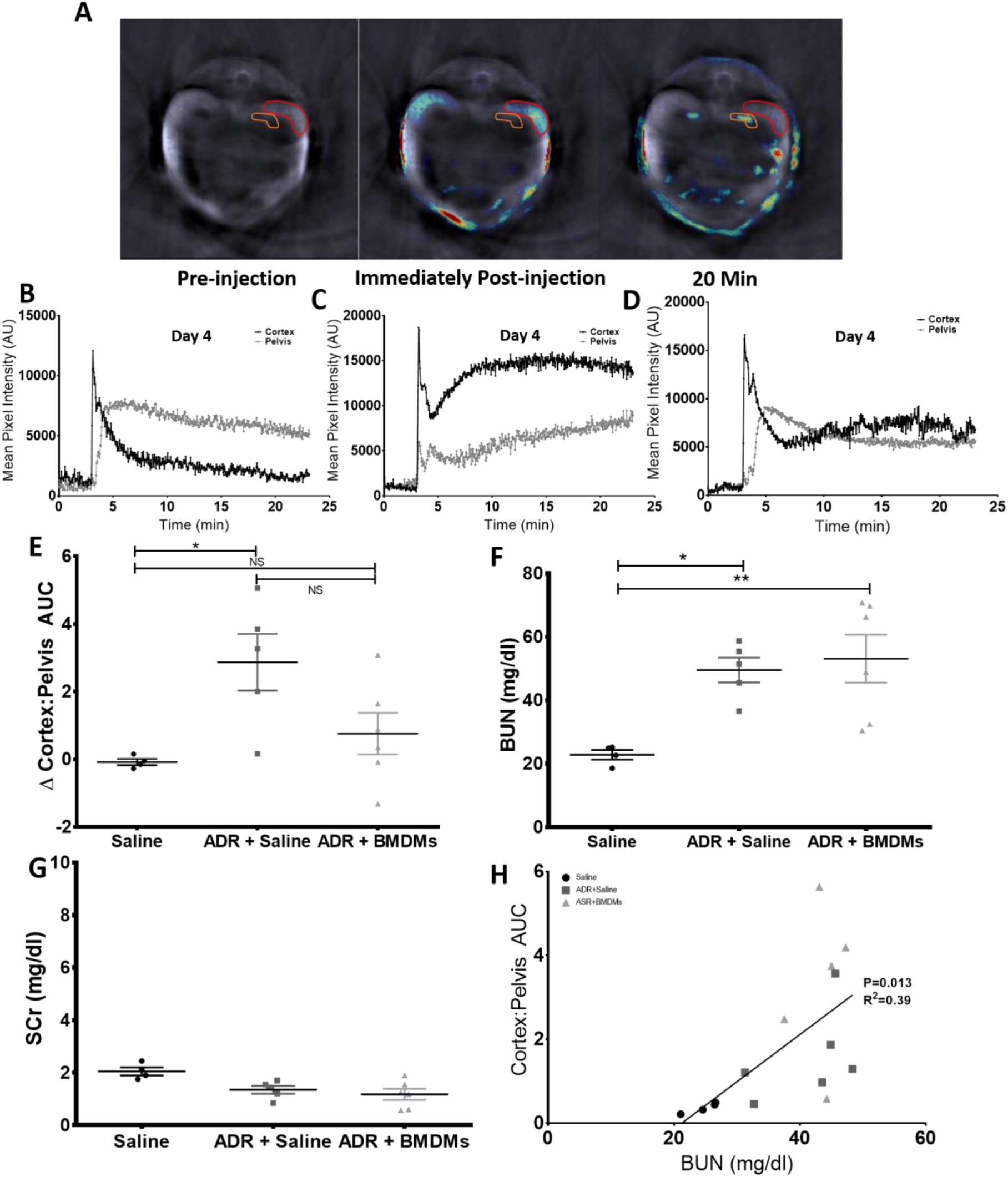
Representative images showing the a typical cross section of a mouse as obtained by the MSOT prior to, immediately post and 20 minutes post IRDye administration, Cortex and medullary regions of interest are shown (A). Clearance kinetics on day 4 of IRDye 800 carboxylate from both the kidney cortex and pelvis regions of interest in a typical healthy (B), ADR treated (C) and ADR+BMDM treated mouse (D). The change in the mean cortex:pelvis AUC between days 1 and 4 in all mice is shown in (E) (Mean ΔAUC C:P −0.085, 0.755 and 2.866). Serum BUN (F) (mean 24.7, 43.4 and 41.1 mg/dl) and SCr (G) levels were quantified on day 4 in each mouse. The correlation between the cortex:pelvis AUC and blood urea nitrogen on day four is shown in (H). Each data point represents an individual mouse. * P<0.05, ** P<0.01

### Use of MSOT to investigate the effect of BMDMs on hepatic function in ADR-dosed mice

We assessed liver function by measuring the clearance of indocyanine green (ICG) which is exclusively eliminated from blood by the liver using MSOT, as previously described (Brillant et al., 2017). Representative MSOT snap shot images illustrate the change in ICG signal in the ischiatic vessel (Fig 4A). Plotting of the signal intensities over time showed that ICG clearance was delayed both in the ADR+BMDM and the ADR groups compared to the saline group (Fig 4 B-D). To investigate the ameliorative potential of BMDMs on liver function, we determined the change in AUC for ICG between days 1 and 4 in each individual mouse (ΔICG AUC) (Fig. 4E). There was a significant elevation in ΔICG AUC in the ADR group compared to healthy controls, but not between controls and ADR+BMDM groups, nor between the ADR and ADR+BMDM groups (Fig. 4E). Alanine aminotransferase (ALT) was significantly elevated in the sera of the ADR and ADR+BMDM groups compared to the controls on day 4 (Fig. 4F), and correlated significantly with ICG AUC (P=0.0009, R^2^=0.59) (Fig. 4G). To investigate the relationship between cardiac, kidney and liver injury, we plotted cardiac output together with the MSOT data for kidney and liver function on a 3-D graph. This shows the relationship between CO, kidney and liver function for each animal (Sup Fig 3).

**Figure 4:**
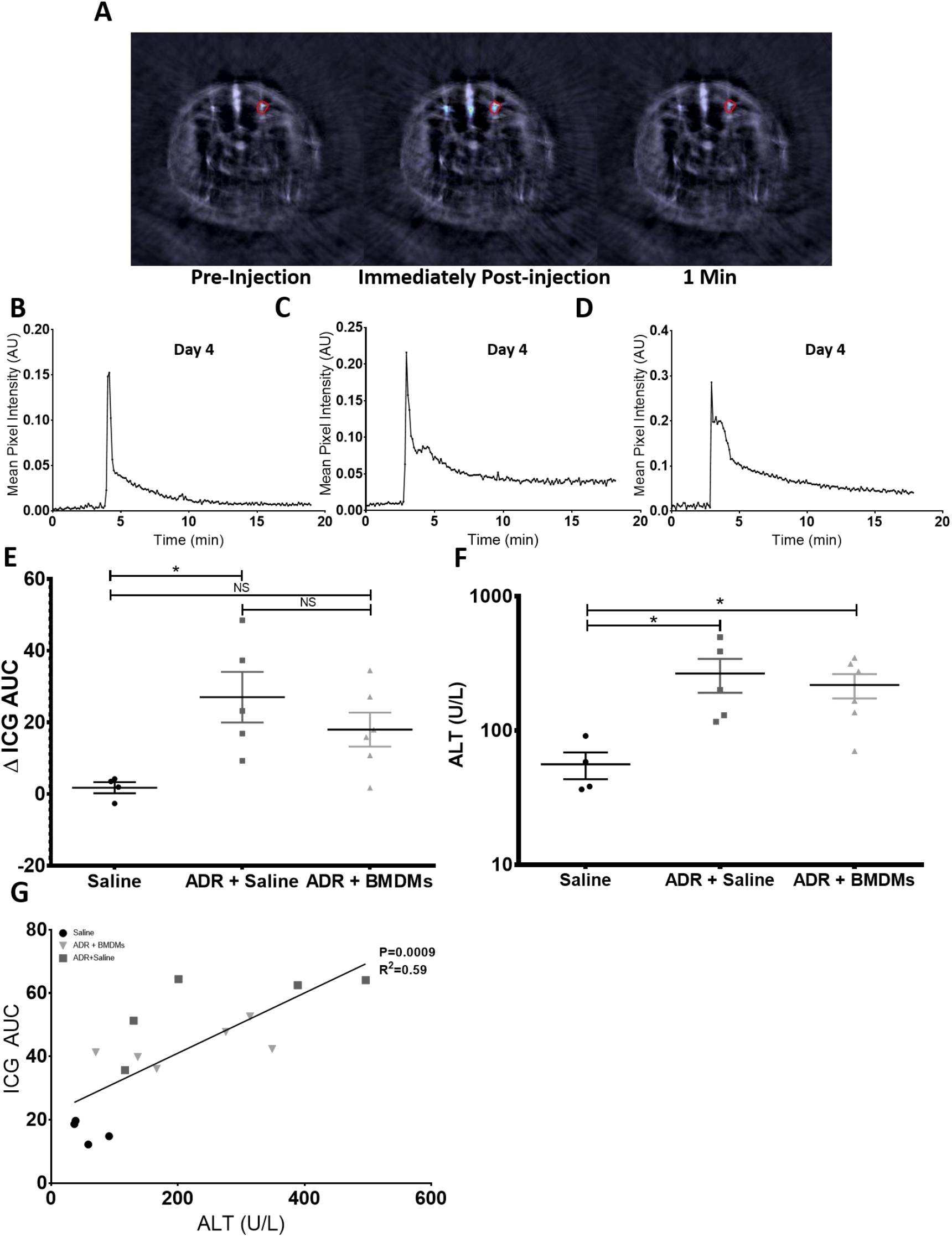
Representative images showing the a typical cross section of a mouse as obtained by the MSOT both prior to, immediately post and 1 minute post ICG administration (A). Clearance kinetics on day 4 of ICG from a typical healthy mouse (B), an ADR treated mouse (C) and an ADR+BMDM mouse (D) from a single ischiatic vessel’s region of interest. Clearance kinetics on day 4 of ICG from a mouse which had received adriamycin from a single ischiatic vessel (C). The change in the mean ICG AUC between days 1 and 4 in all mice is shown in (E) (Mean ΔAUC ICG 1.8, ΔAUC ICG 18). Serum ALT was quantified in all mice on day 4 (F). The correlation between the ICG AUC and alanine aminotransferase on day four is shown in (G) (mean 56.1, 218.6 and 266.6 U/L). Each data point represents an individual mouse. * P<0.05.

### BMDMs failed to ameliorate ADR-induced histological damage in the kidney and liver

The functional data from the MSOT analyses suggested that BMDMs had a tendency to improve renal and hepatic function in ADR-dosed mice, but did not improve cardiac function. To investigate whether the apparent improvement in renal and hepatic function was associated with an amelioration of tissue damage, histological analysis of the liver and kidneys of mice was carried out at the study endpoint. Kidney sections were assessed for the presence of intratubular protein casts and flattened tubular epithelium, and liver sections for hepatocellular degeneration and necrosis. There was a significant difference between healthy and ADR injured animals, but in contrast to the functional data, administration of BMDMs failed to improve the extent of histological damage in the kidney or liver. Histologically, the kidneys of control mice showed no evidence of injury (Fig 5A) whereas kidneys of ADR mice had intratubular protein casts and flattening of tubular epithelial cells (Fig 5B, arrow) regardless of BMDM administration. Likewise, the livers of control mice showed no signs of injury (Fig 5C) whereas ADR and ADR+BMDM mice showed evidence of hepatocellular degeneration (Fig 5(1)) and necrosis (Fig 5D(2)). Histological scoring in both the kidney and liver showed that control mice had no evidence of histological damage whereas both the ADR and ADR+BMDM groups showed varying degrees of injury (Fig 5E, F respectively).

**Figure 5:**
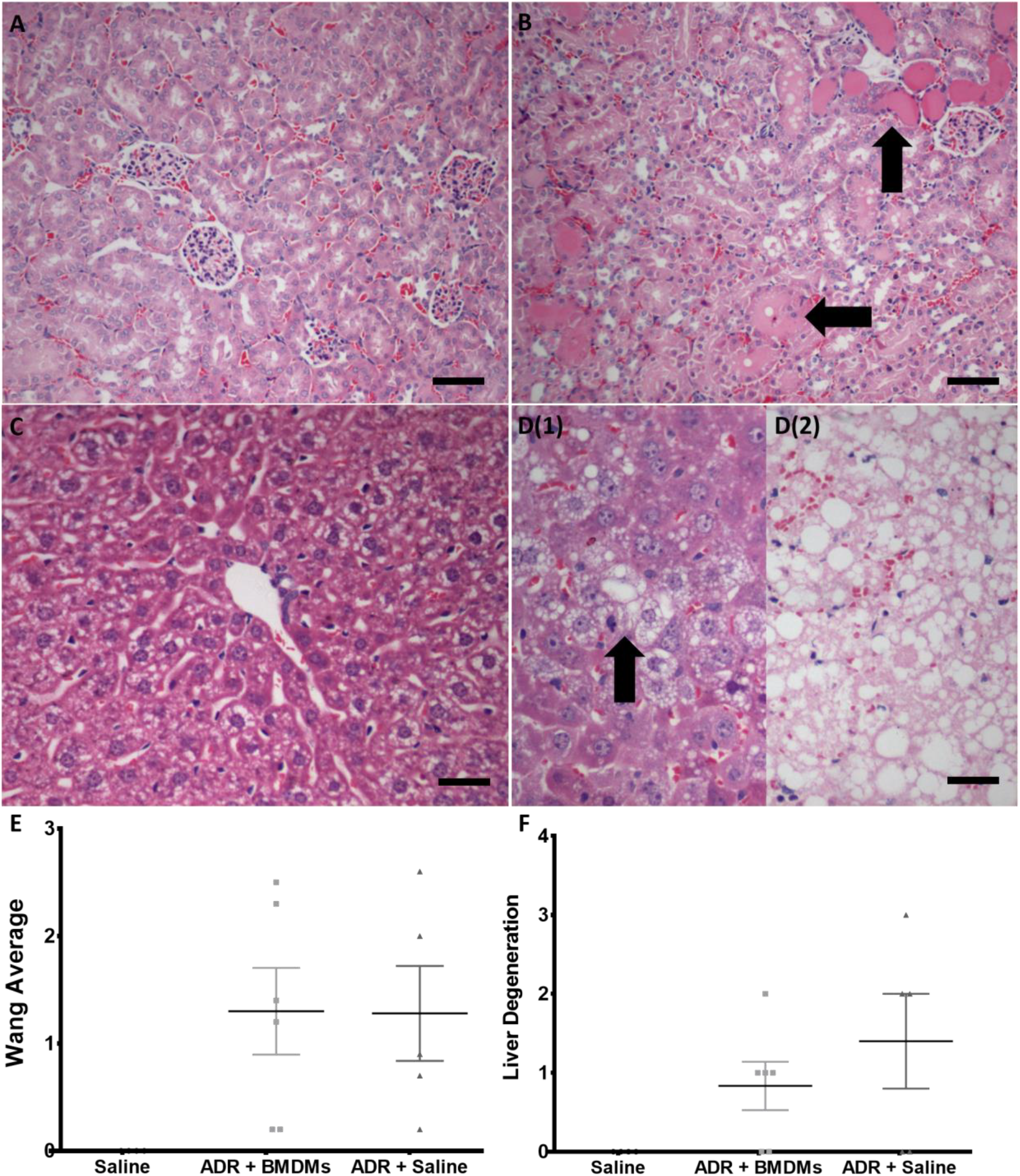
Histological analyses from both the kidney and liver. Histological analyses show sections from typical sections of a kidney from a healthy mouse (A), the kidney of a mouse which had received adriamycin (B), the liver of a healthy mouse (C) and the liver of a mouse which had received adriamycin (D1, 2). All organs were collected on day 4 of the study and were stained with haematoxylin and eosin. (A) and (B); scale bar = 100 μm. (C) and (D); scale bar = 50 μm. No evidence of injury are observed in (A) or (C). (B) shows evidence of intratubular protein casts and flattening of the tubular epithelium (arrow). (D1) shows evidence of hepatocellular degeneration (arrow) and (D2) shows evidence of necrosis with evidence of pale eosinophilic substance. Histological scoring for the kidney and liver are shown (E, F respectively). Each data point represents an individual animal.

### Bioluminescence imaging shows that BMDMs accumulated in the kidneys and liver following ADR-induced injury, but not in the heart

In order to investigate the effect of organ injury on BMDM distribution, mice were imaged on the same day, or 3 days after administration of luciferase+ BMDMs, which corresponded to the 1^st^ and 4^th^ day following saline or ADR administration. Bioluminescence imaging showed that on day 1, cells were mostly in the lungs of both control and ADR animals. By day 4, BMDMs were no longer detectable in the controls, but had a widespread distribution in the ADR group (Fig 6A, B). Given the poor spatial resolution of bioluminescence imaging, it was not possible to determine which organs the BMDMs had populated in the ADR group at day 4. Therefore, immediately after in vivo imaging on day 1, three animals were immediately sacrificed to quantify the biodistribution of BMDMs in the major organs ex vivo, with the remaining three mice being sacrificed on day 4. Similarly to the in vivo data, ex vivo analysis of organs on day 1 showed no obvious difference between control and ADR mice, with most BMDMs being in the lungs, and some detected in the spleen (Fig. 6C, D). By day 4, BMDMs could only be detected in the lungs of control animals, but were present in the lungs, spleen, kidneys and liver of ADR animals. BMDMs were not detected in the hearts of control or ADR animals (Fig. 6E, F).

**Figure 6:**
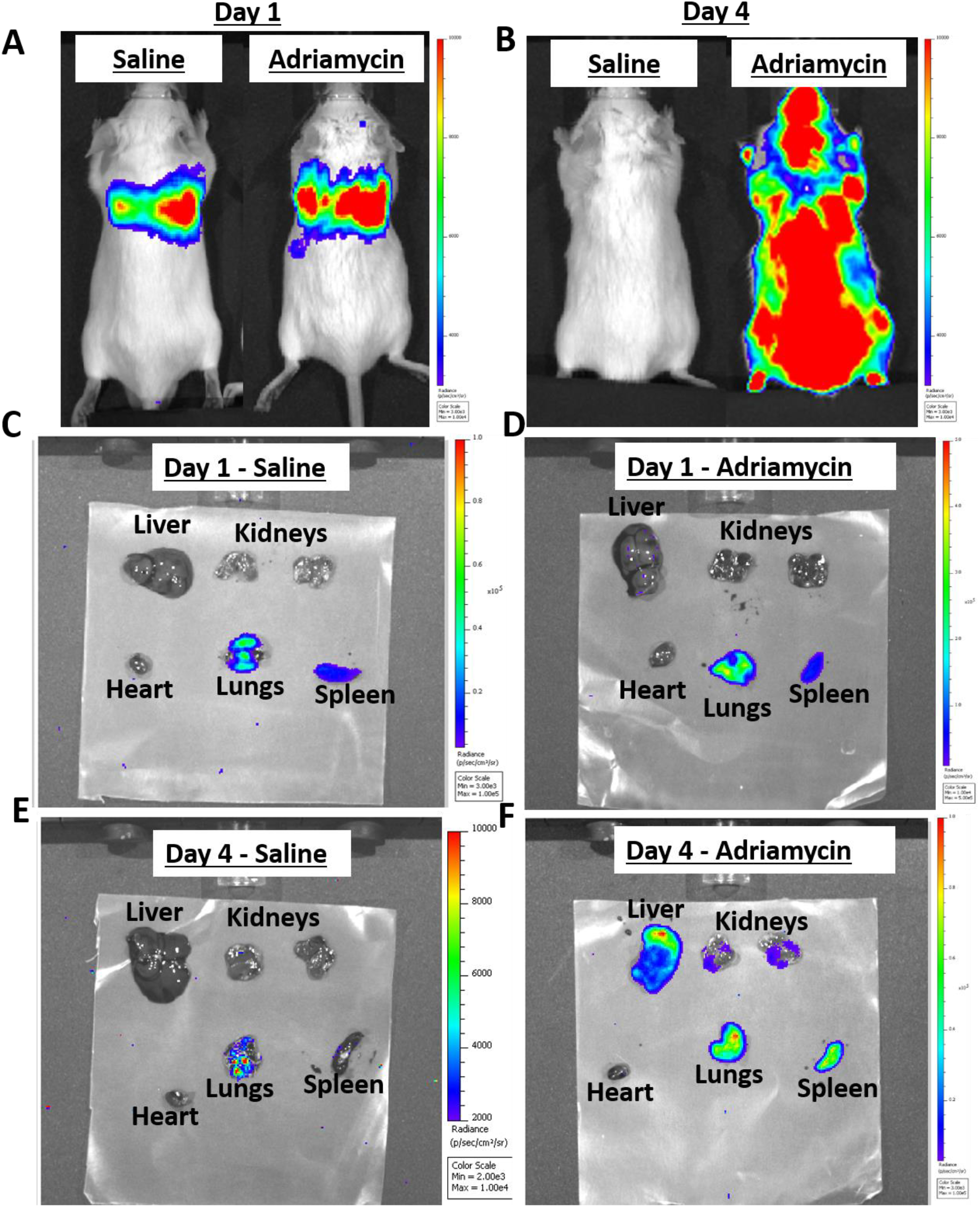
All mice received 10^7^ primary bone marrow derived macrophages which expressed luciferase via tail vein administration. Whole body images of typical mice treated with either saline or Adriamycin on both days 1 (A) and 4 (B). Ex vivo bioluminescence images from typical saline (C, E) and Adriamycin (D, F) treated mice on both days 1 (C, D) and 4 (E, F). Individual scales are shown on the right of each image. Each image shows the liver, kidneys, heart, lungs and spleen from an individual mouse. Scales for all ex vivo images are identical, as are scales for all whole body images.

We measured the total flux from the individual organs ex vivo to quantify the differences in organ biodistribution between the treatment groups and time points (Fig. 7). The total flux from the heart, lungs and spleen decreased between days 1 and 4 in both the control and ADR mice (Fig 7A, B, C). The same trend was observed for the liver and kidney in the controls, with total flux decreasing between days 1 and 4. On the other hand, in the ADR mice, total flux increased significantly in the liver and kidneys between days 1 and 4 (Fig 7D, E).

**Figure 7:**
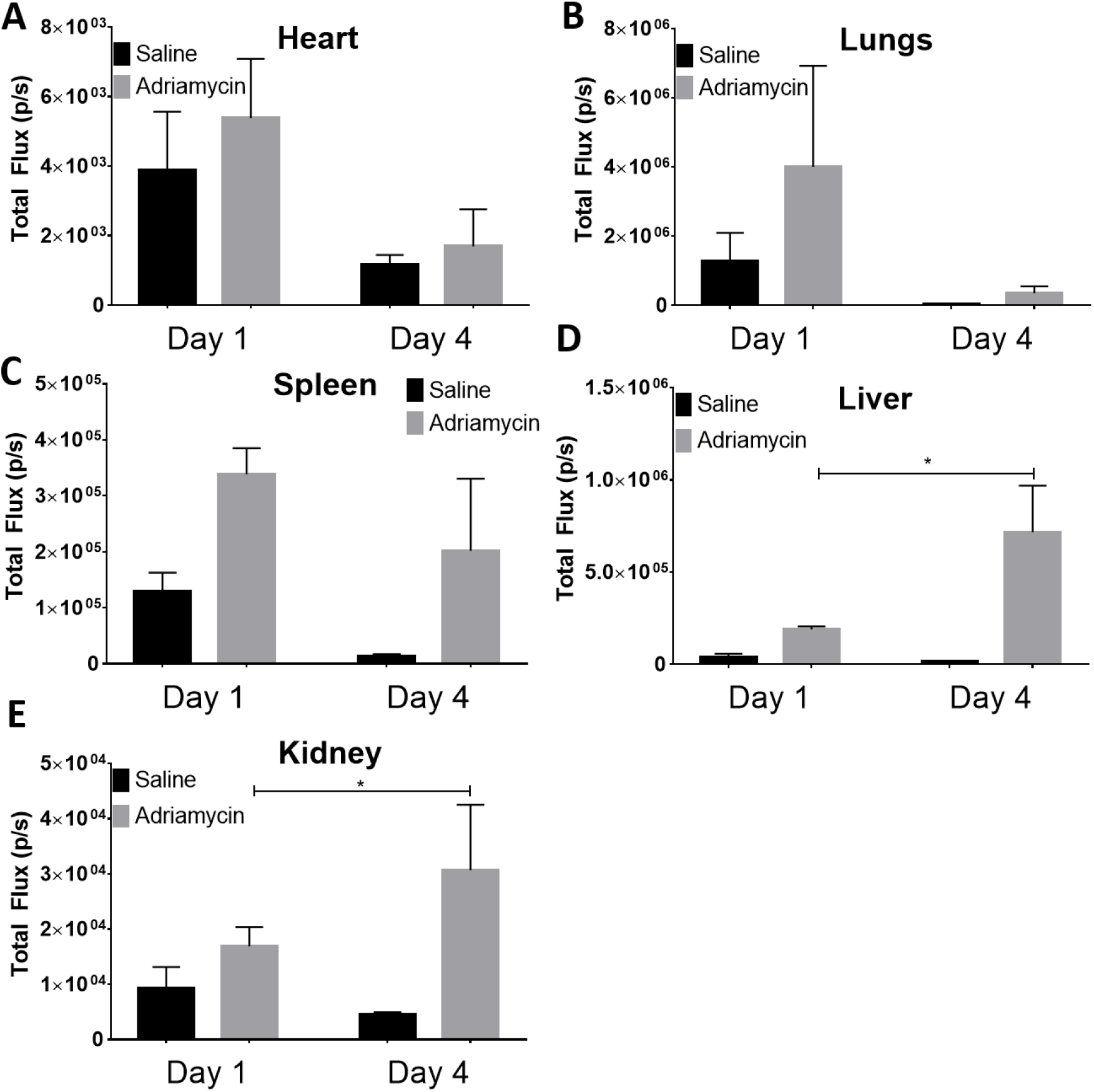
Quantified ex vivo bioluminescent results from organs of mice that received either saline on Adriamycin on both days 1 and 4. Data is shown for the heart (A), the lungs (B), the spleen (C), the liver (D) and the mean of both kidneys (E). Each bar represents the mean of 3 individual animals. * P<0.05

## Discussion

In the current investigation, cardiac function was measured by ultrasonography alongside hepatic and renal function by MSOT under the same anaesthesia session. This allowed us to determine the functions of 3 separate organs utilising a multimodal imaging approach. The cardiac measurements taken easily and quickly with the ultrasound software revealed that there were no significant changes in fractional shortening or ejection fraction between days 1 and 4 in the ADR and ADR+BMDM groups when compared to the healthy mice. However, significant decreases in cardiac output and stroke volume between days 1 and 4 where detected when comparing the saline treated mice to those which received adriamycin, regardless of the addition of the RMT. This suggests that a single dose of adriamycin significantly reduces cardiac function between days 1 and 4 of this study; however, the addition of BMDMs as a RMT show no beneficial effect on cardiac function.

We have previously used MSOT to monitor renal function in a mouse model of adriamycin-induced chronic kidney disease by measuring the clearance kinetics of IRDye (Scarfe et al., 2015). The two parameters used in this earlier study, which focussed on the chronic stage of renal injury, were the Tmax delay, and the exponential decay time of the IRDye in the renal cortex, which were measured at week 5 following ADR dosing, when the nadir of acute renal injury had passed. However, in the current study, which focussed on the acute phase of adriamycin-induced injury, these parameters were not appropriate because many animals did not display an IRDye ‘peak’ in the pelvis, nor an exponential decay in the renal cortex. Therefore, we considered that the most appropriate parameter in this case would be the ratio of the area under the curve (AUC) of the cortex and pelvis kinetic curves (AUC C:P), as this measurement reflects both impaired clearance through the cortex and delayed accumulation in the pelvis, and could be measured in all animals. A single dose of adriamycin caused a significant reduction in kidney function when compared to healthy mice over a 4 day period. Administration of M2 BMDMs 1 day after adriamycin administration resulted in a non-significant reduction in kidney function when compared to control mice. The AUC C:P showed a strong positive correlation with a more traditional biomarker of kidney injury, BUN, which reinforces the utility of AUC C:P as a measure of kidney function. There were, however, no differences between the mean BUN measurements in mice which received adriamycin alone and those which received adriamycin and M2 BMDMs. SCr was not significantly different between any of the groups, which highlights the lack of sensitivity of this biomarker for indicating renal function in rodents. Histological analyses of the kidneys showed evidence of intratubular protein casts and flattening of tubular epithelium in mice which received adriamycin, irrespective of whether they were administered BMDMs or not. Therefore, although MSOT showed that the M2 BMDMs appeared to cause a subtle improvement in renal function, there was no corresponding improvement in histological damage or BUN levels. A study by Lu and colleagues had previously demonstrated that M2 macrophages reduce inflammatory infiltrates in adriamycin-induced nephropathy; however, macrophages were not administered until 5 days after adriamycin administration and biomarker analyses not carried out until day 28 (Lu et al., 2013). It is possible in this current study that analysis of kidney function at later time points than day 4 may have yielded more significant improvements in function. Wang and colleagues studied the effect of M0, M1 and M2 activated macrophages in a mouse model of chronic adriamycin nephropathy (Wang et al., 2007). These authors had found that by 4 weeks, M0 macrophages had no effect on renal injury, while M1 macrophages promoted histological damage, and M2 macrophages significantly ameliorated tubular and glomerular injury (Wang et al., 2007). By contrast, we failed to observe any ameliorative effects of the M2 macrophages on tissue damage, but this may have been because our analysis was performed at 4 days, rather than at 4 weeks. Consistent with our current study, Wang and colleagues showed that M2 macrophages trafficked to inflamed kidneys. In addition, they provided evidence that the exogenous M2 macrophages ameliorate injury by reducing the infiltration of resident macrophages, thereby reducing inflammation.

In the current study the clearance of ICG from the blood of mice was used as a measure of liver function. ICG is a fluorescent cyanine dye which is used clinically in a number of diagnostic procedures including measurement of cardiac output, liver blood flow, ophthalmic angiography and hepatic function (Caesar et al., 1961, Okochi et al., 2002,Iijima et al., 1997). It absorbs in the near infrared region which makes it an ideal optoacoustic contrast agent. This, combined with the fact that ICG is microsomally metabolised in the liver and cleared through the hepatobiliary route make it an ideal agent for determination of liver function with MSOT. Using the ICG AUC we show here that there was no change in liver function in healthy mice between days 1 and 4 whereas mice which received adriamycin had a significantly decreased liver function in the same time window. Liver function was also decreased between days 1 and 4 in mice which received adriamycin and M2 BMDMs but not significantly so when compared to healthy mice. This suggests that a single high dose of adriamycin reduces liver function while the addition of M2 BMDMs has a beneficial effect on the function of the liver. The ICG AUC on day 4 of the study correlates strongly and significantly with serum ALT levels, reinforcing the utility of ICG AUC as a functional liver parameter. Histological analyses showed no evidence of liver injury in healthy mice while hepatocellular degeneration and necrosis was observed in all mice which received adriamycin. Again, much like the kidney histopathology and biomarker analysis, there were no differences in ALT or liver histology between mice which received adriamycin alone and those which received adriamycin and the M2 BMDMs. Previous work by Thomas and colleagues found that exogenous unpolarised BMDMs had a beneficial effect on mice with carbon tetrachloride induced liver injury through the recruitment of MMP (matrix metalloproteinase)- producing host cells into the liver (Thomas et al., 2011). This in turn increased host macrophage recruitment and elevated IL-10 and MMP levels. The study also showed modest but significant increases in serum albumin, suggesting liver regeneration (Thomas et al., 2011). The results we present here also suggest a modest improvement in liver function as measured by MSOT, albeit insignificantly. It should be noted that the effect of M2 BMDMs on adriamycin-induced liver injury has not been previously investigated.

The aforementioned multimodal imaging strategy can monitor changes in the functions of the liver, kidney and heart, and identify potential efficacy of M2 BMDMs as a RMT. To investigate the relationship between biodistribution of the M2 BMDMs and any therapeutic effect, we administered M2 BMDMs expressing firefly luciferase which allowed the luminescent imaging and quantification of cells in the mice and organs (three on day 1 and three on day 4). At day 1, we observed no difference in luminescence distribution between both groups of mice, with signals only present in the lungs and spleen after IV cell administration. This pattern of biodistribution has been reported widely after IV administration of cells (Sharkey et al., 2016). However, on day 4 of the study the biodistribution of the cells in saline or adriamycin-dosed mice had changed dramatically. In the saline treated mice there remained only a weak luminescent signal in the lungs of the mice, indicating the cells rapidly die after IV administration. However, in mice which received adriamycin this was not the case; luminescence was detected over the whole body of the mouse when imaged *in vivo,* and in the liver, kidneys, lungs and spleen, but not the heart, when imaged *ex vivo*. While the luminescent signal intensity decreased from day 1 to 4 in the lungs and spleen, it was still observable. On the other hand, the signal intensity in the liver and kidneys was significantly elevated between days 1 and 4. Of note, the total flux in the whole body images on day 4 in ADR treated mice appeared greater than on day 1; however M2 BMDMs do not have a great proliferative capacity so this was likely a result of the weight loss in the ADR treated mice causing less attenuation of light.

### Summary

This investigation demonstrates that changes in the function of the liver, kidney and heart can be tracked over time in individual healthy animals and those that have received a single high dose of adriamycin. We show that these changes in liver and kidney function correlate well with traditional serum biomarkers of injury and show histological evidence of injury. We also show that the addition of M2 BMDMs improves kidney and liver function over the study as measured by MSOT. However neither biomarker nor histological analyses showed a reduction in the severity of injury. This may suggest that subtle changes in organ function can be detected using MSOT imaging prior to changes in organ histopathology and accumulation of serum biomarkers. Therefore, MSOT allows the assessment of the efficacy of a potential RMT in mice without the need for repeated blood or urine sampling. Secondly this study demonstrates an imaging technique to monitor the biodistribution of M2 BMDMs in healthy animals and those with organ dysfunction. Unfortunately, to accurately assess the intra-organ biodistribution of M2 BMDMs, it is required that animals are sacrificed prior to imaging. We show that adriamycin affected the biodistribution of M2 BMDMs since mice with kidney and liver dysfunction demonstrated an increase in longevity and migration of the therapeutic cells to both organs, which incidentally also appeared to provided signs of efficacy in the functional study. No signs of efficacy were observed in the heart, which showed no evidence that BMDMs migrated towards it.

This current study describes and demonstrates an “imaging toolbox” to assess murine renal, hepatic and cardiac function in a minimally invasive manner using both MSOT and ultrasound in a single anaesthesia session. Comorbidities are common clinically so the ability to utilise an imaging toolbox to investigate the potential of regenerative therapies in preclinical species is crucial. This toolbox allows more accurate assessment of the efficacy of potential regenerative therapies than current histological and biomarker analyses as it allows the extent of the recovery to be measured in individual animals over time. Many regenerative therapies which show efficacy in preclinical species are not effective clinically which may be a result of improper methods to assess their efficacy in individual animals preclinically. This toolbox may allow more accurate assessment of efficacy and enable a more robust determination of the risk: benefit ratio of a potential regenerative therapy prior to clinical translation and therefore reduce the number of therapies that don’t show true efficacy that are tested in human patients. This toolbox also improves understanding of the mechanism in which cell therapies elicit efficacy through the study of their biodistribution. We can determine whether cells must reach the target organ and engraft to show efficacy, which is important for determining the optimal administration route.

## Materials and Methods

### Animals

Mice were purchased from Charles River, UK, and were housed with *ad libitum* access to food and water. All animal experiments were performed under a license granted under the Animals (Scientific Procedures) Act 1986 and were approved by the University of Liverpool ethics committee. Experiments are reported in line with the ARRRIVE guidelines.

### Primary macrophage isolation

Primary bone marrow derived macrophages (BMDMs) were prepared as previously described (Sharkey et al., 2017). Male BALB/c mice were used to isolate BMDMs for the efficacy study whereas for the biodistribution study, mice with a mixed background (L2G85 mice bred to wild type FVB mice) expressing the CAG-luc-eGFP L2G85 transgene were used (FVB-Tg(CAG-luc,-GFP)L2G85Chco/J). Briefly, femurs and tibias of mice (8-10 weeks) were harvested and muscle tissue removed from the bones in a sterile fume hood. Bone marrow was flushed from the bones using a sterile syringe with Dulbecco’s Modified Eagle’s Medium (DMEM): F12 cell culture medium (Gibco) supplemented with 10 % foetal bovine serum 2mM glutamine and 1 x penicillin/streptomycin (Invitrogen). The bone marrow was suspended in the medium before being passed through a cell strainer (40 μm) and then cultured in DMEM: F12 media containing 20 ng/ml murine recombinant macrophage colony stimulating factor (MCSF-1). Bone marrow suspensions were cultured at 37 °C, 5 % CO_2_ and medium was replaced every other day. On day 7, macrophages were considered fully differentiated as determined by the expression of both CD11b and F4/80 (Biolegend) by flow cytometry. Mature BMDMs were then polarised towards an M2-like phenotype by the overnight addition of recombinant murine interleukin (IL)-4 (20 ng/ml).

### Induction of organ dysfunction

Male BALB/c mice (8 – 10 weeks) received either adriamycin (20 mg/kg, n=11) or saline (0.9 %, n=4) intra-peritoneally (IP) on day 0. Adriamycin (doxorubicin hydrochloride, Tocris Bioscience) was dissolved in warm saline (0.9 %) to make a stock solution (10 mg/ml) before administration. Mice were weighed on a daily basis to monitor their wellbeing and mice which received adriamycin were provided with a wet food diet.

### Imaging protocol

Imaging was carried out on day 1 and 4 under the same imaging session and using the following protocol: Mice were anaesthetised and fur was removed from the torso by shaving and epilating. Mice were then imaged by ultrasound to generate functional cardiac parameters. The tail veins of mice were then cannulated and mice moved to the MSOT system and received ICG (Carl Roth, Germany) for liver functional measurements. The catheter was flushed with saline before administration of IRDye800 carboxylate (LI-COR) for functional liver measurements. Mice were then allowed to recover in a heat box before being returned to their home cage. A schematic showing how the study was carried out can be found in Fig. 1. Results for MSOT and ultrasound analyses are expressed as the change in each parameter between days 1 and 4 in the study. Detailed descriptions of each imaging protocol are provided below.

### Assessment of cardiac function

Cardiac function was assessed using the Prospect 2.0 ultrasound system (S-Sharp, Taiwan). Mice were anaesthetised using isofluorane and oxygen and the mice were placed dorsally on a heated platform. Mice were fixed in place during imaging using surgical tape. Ultrasound gel was applied to the chest area of the mice and the ultrasound transducer positioned above the chest area. The following parameters were measured: epicardial area and endocardial area in the long axis view, left ventricle length, epicardial areas in the short axis view, M-mode images of both the long and short axis views in order to measure heart rate, left ventricular interior diameter and wall thickness in both diastole and systole. These parameters were used to calculate fractional shortening (FS), ejection fraction (EF), stroke volume (SV) and cardiac output (CO).

### Assessment of liver function

Liver function was assessed using the inVision 256-TF MSOT imaging system (iThera Medical, Munich). Immediately after assessment of cardiac function the mice were moved to the MSOT imaging system. Prior to being placed into the system the tail vein of the mice was cannulated to allow injection of the optical imaging contrast agents during photoacoustic imaging. Mice were placed in the system and allowed to acclimatise for 15 minutes prior to recording data. Imaging focussed on the ischiatic vessels close to the hips of the mice for detection of indocyanine green (ICG) and its subsequent clearance. The following parameters were used: an acquisition rate of 10 frames per second (consecutive frames averaged to minimalise effects of respiration movement), with wavelengths of 700, 730, 760, 800, 850 and 900 nm being recorded. Data were recorded for 3 minutes prior to intravenous (IV) injection of ICG (40 nmol, 100 μl) over a 10 s period. Data were reconstructed using a model linear algorithm (View MSOT software) and multispectral processing using linear regression for ICG, deoxy- and oxy- haemoglobin spectra to resolve the signal for the ICG dye. Regions of interest drawn around the ischiatic vessels of each mouse were used to quantify the ICG dye signal (as mean pixel intensity) in the vessels of the mice. These data were used to calculate the area under the clearance curve (AUC). Data expressed as the change in ICG AUC in each individual mouse between days 1 and 4 (ΔICG AUC).

### Assessment of kidney function

Kidney function was assessed utilising a similar method as the assessment of liver function. After liver function assessment, the catheter was flushed with a small amount of saline. Data were recorded using the following parameters: wavelengths of 775 and 850 nm and an acquisition rate of 10 frames per second (consecutive frames averaged). Imaging focussed on the centre of the right kidney of the mouse where the renal pelvis was visible. Data was recorded for 3 minutes prior to the injection through the tail vein catheter of IRDye 800 carboxylate (20 nmol, 100 μl) over a period of 10 s. Data was reconstructed using a model linear algorithm and a difference protocol (775 nm – 850 nm) to resolve the signal for the IRDye 800 carboxylate. Regions of interest were drawn around the renal cortex and the renal papilla/pelvis region of the right kidney in each mouse to quantify the IRDye signal in the kidney. The mean pixel intensity data were used to calculate the AUC of both the renal cortex and papilla/pelvis. Data were expressed as the change in the ratio between the AUC of the cortex and the AUC of the pelvis regions between days 1 and 4 (ΔAUC C:P).

### Therapeutic cell administration

M2-like primary BMDMs were administered to mice which received adriamycin (n=6) on day 1 immediately after cardiac, hepatic and renal imaging. BMDMs were harvested from low adherence flasks (Corning) after maturation and polarisation by gentle agitation and scraping. Cells were then counted and suspended to a concentration of 10^7^ cells/100 μl saline. After mice were removed from the MSOT imaging system, 100 μl of the cell suspension was administered via the tail vein cannula and mice were allowed to recover in a heat box before being returned to their home cage. Mice which did not receive BMDMs received 100 μl saline via the tail vein catheter.

### Quantification of serum biomarkers

On day 4 mice were culled using an increasing concentration of CO_2_ and exsanguinated by cardiac puncture. Blood was allowed to clot at room temperature before centrifugation to isolate the serum.

Blood urea nitrogen (BUN, QuantiChrom Urea Assay Kit, BioAssay Systems), serum creatinine (SCr, Serum Creatinine detection kit, ARBOR ASSAYS) and alanine aminotransferase activity (ALT, Thermo Fisher) were quantified according to manufacturer’s instructions in a 96 well plate and were read using a FLUOstar Omega microplate reader (BMG LABTECH).

### Histopathological analysis

Kidney and liver tissues were fixed in 4 % paraformaldehyde (4 °C) for 24 hours before being washed in PBS, dehydrated with increasing concentrations of ethanol and subsequently embedded in paraffin. The kidneys were sectioned in order to obtain two symmetrical halves, to include the cortex, and medulla extending to the renal pelvis. For the liver, the large lobe was collected for anlysis. Tissue sections were cut (5 μm) and were stained with haemotoxylin and eosin (HE) and Periodic Acid Shiff (PAS) by standard methods. Kidney injury was scored 0-5 on 10 consecutive 200x microscopic fields in the outer stripe of the outer medulla and cortex on PAS stained slides adapting the method described from Wang and colleagues (Wang et al., 2005). For liver, histological sections were assessed semi quantitatively for degeneration and necrosis following a semi quantitative scale representative for the section area involved (0=0%; 1=1-25%; 26-50%; 51-75%; 76-100%). Veterinary pathologist (LR) was blinded in regard to the experimental groups.

### Determining cellular biodistribution

To determine cellular biodistribution by bioluminescence imaging, male BALB/c mice (8 – 10 weeks) were used. The same adriamycin and macrophage dosing schedule was used as in the previous study: Mice received either 20 mg/kg adriamycin IP (n=6) or saline IP (n=6). 24 hours later all mice received IV injections of 10^7^ PMDMs isolated from mice expressing the CAG-luc-eGFP L2G85 transgene. All mice received luciferin (1.5 mg/kg, IP) before being imaged using an IVIS Spectrum *in Vivo* Imaging System (Perkin Elmer). Mice were imaged in both dorsal and ventral positions using an automatic exposure time before being sacrificed. Organs were then dissected and imaged using the same protocol. The remaining mice were returned to their home cages until day 4 post adriamycin administration (the study end point) when they were imaged as previously described.

### Statistical analysis

Statistical analyses were carried out using Origin software. One-way ANOVA analysis was used for comparison of two groups and one-way ANOVA followed by Tukey analysis was used to compare multiple groups. Results were determined to be significant when P < 0.05. To assess whether correlations between results were significant Pearson’s correlation coefficient was calculated.

## Acknowledgements

This work was supported by the UK Regenerative Medicine Platform Safety and Efficacy Hub (grant ref MR/K026739/1). The authors report no competing interests.

---

**Supplementary figure 1:**
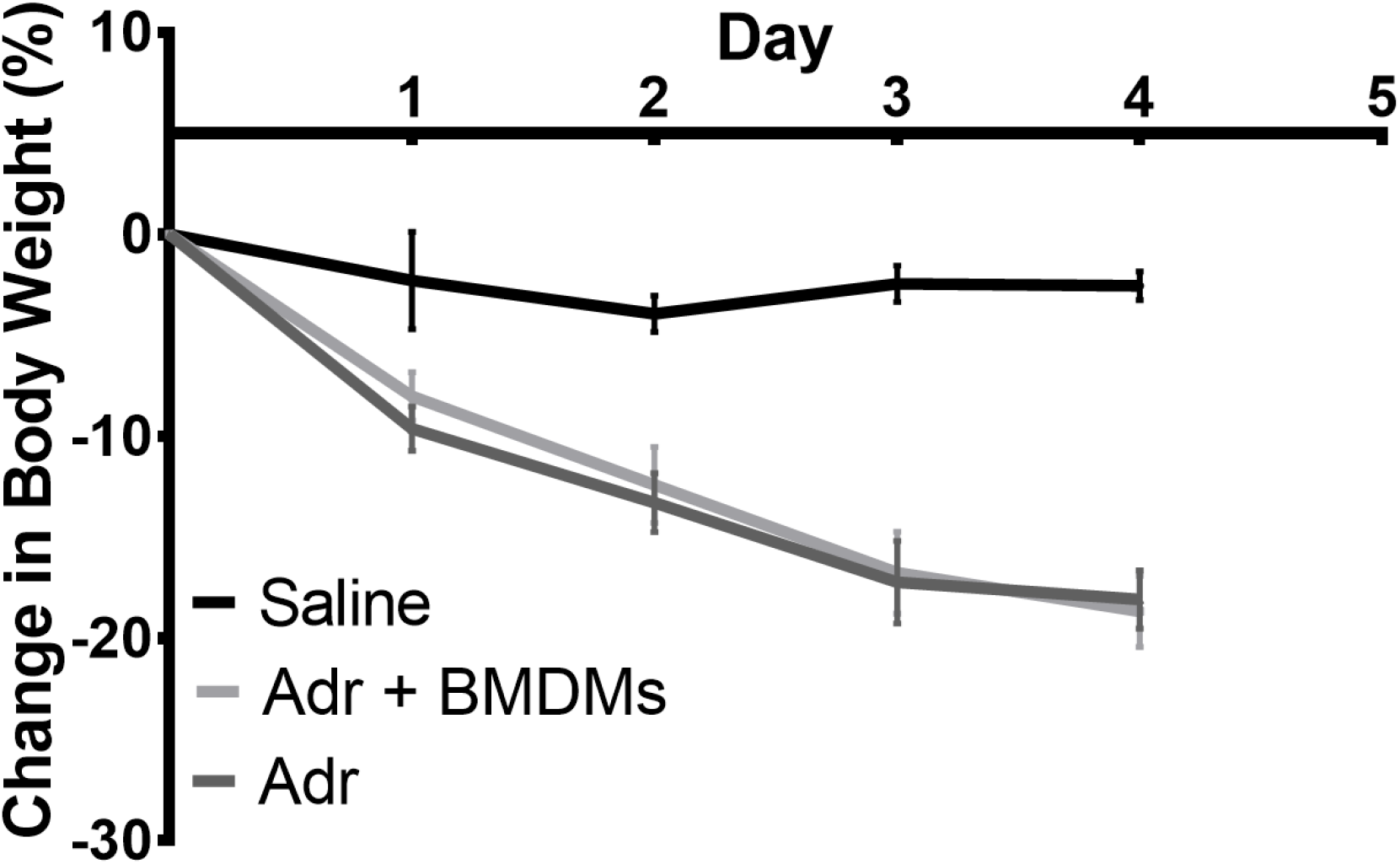
Change in body weight in mice which received saline, ADR+BMDMs and ADR. Each data point represents the mean of mice within the group

**Supplementary figure 2:**
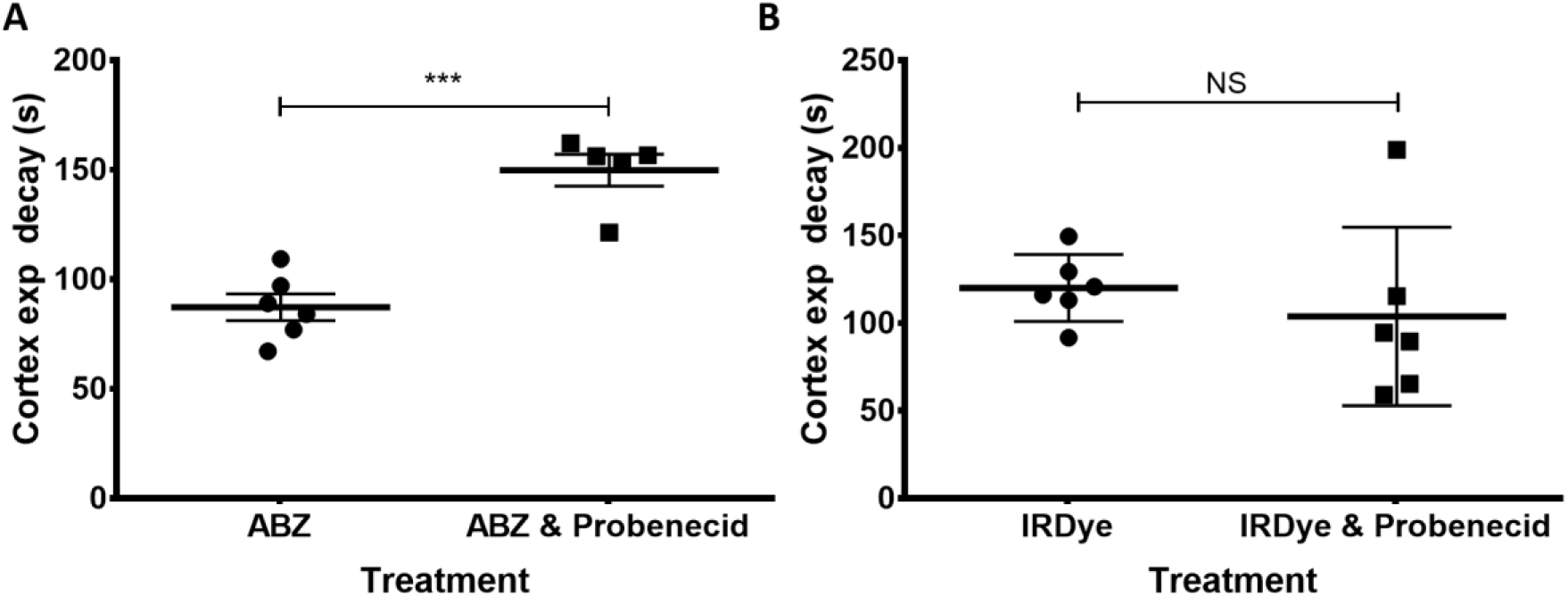
The effect of 50 mg/kg Probenecid on the clearance of two near infrared dyes through the cortex of the kidneys of mice as measured by MSOT. Results are expressed as the exponential decay time (s) of each dye from the cortex of the mice. Each data point represents an individual animal. NS = non-significant, *** P=0.001.

**Supplementary Figure 3:**
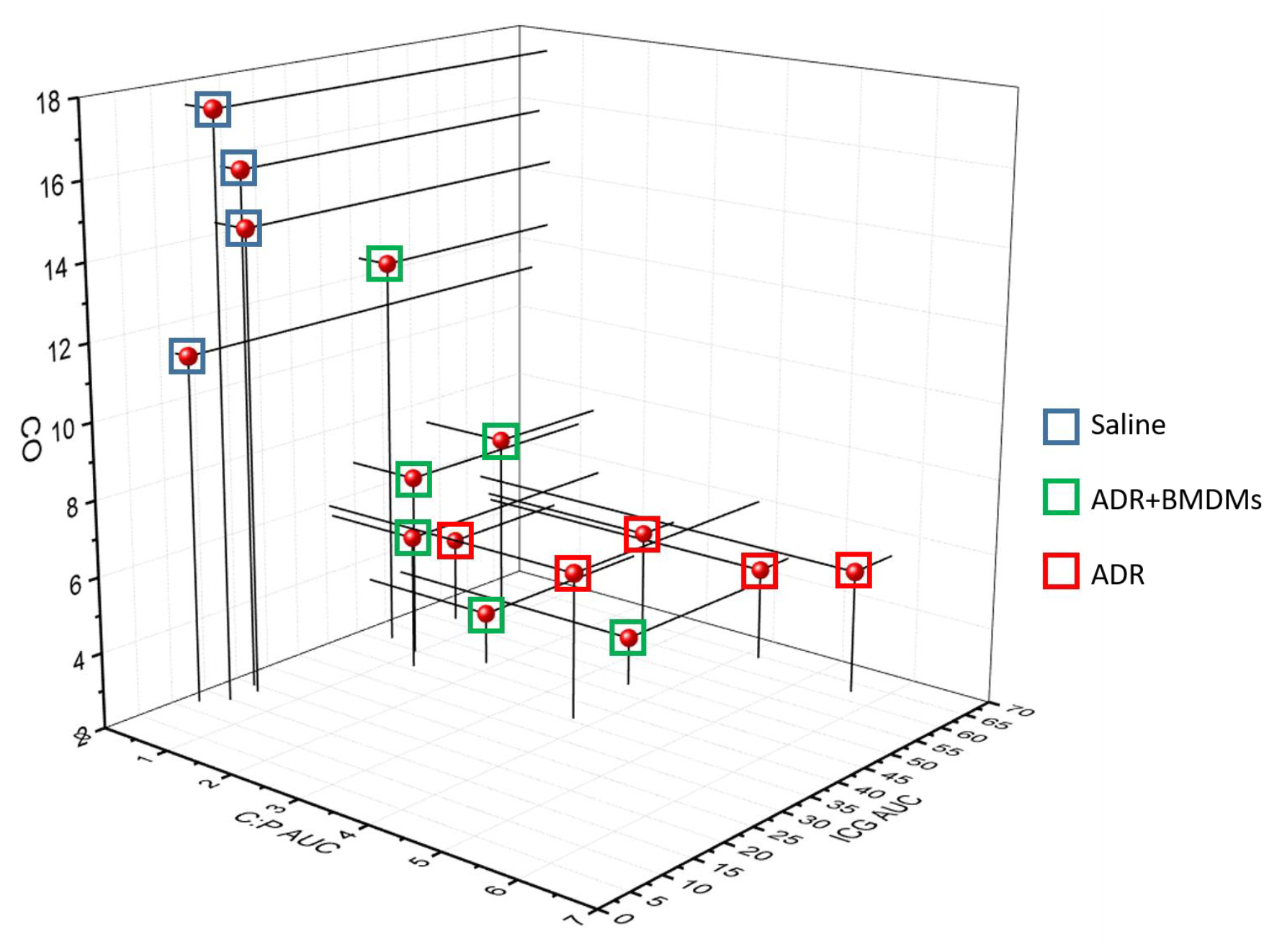
3D dot plot showing the relationship between the change in cardiac output, C:P AUC and the ICG AUC between days 1 and 4 in saline treated mice (blue boxes), ADR treated mice (red boxes and mice treated with ADR and BMDMs (green boxes). Each dot represents an individual mouse

**Supplementary Table 1:**
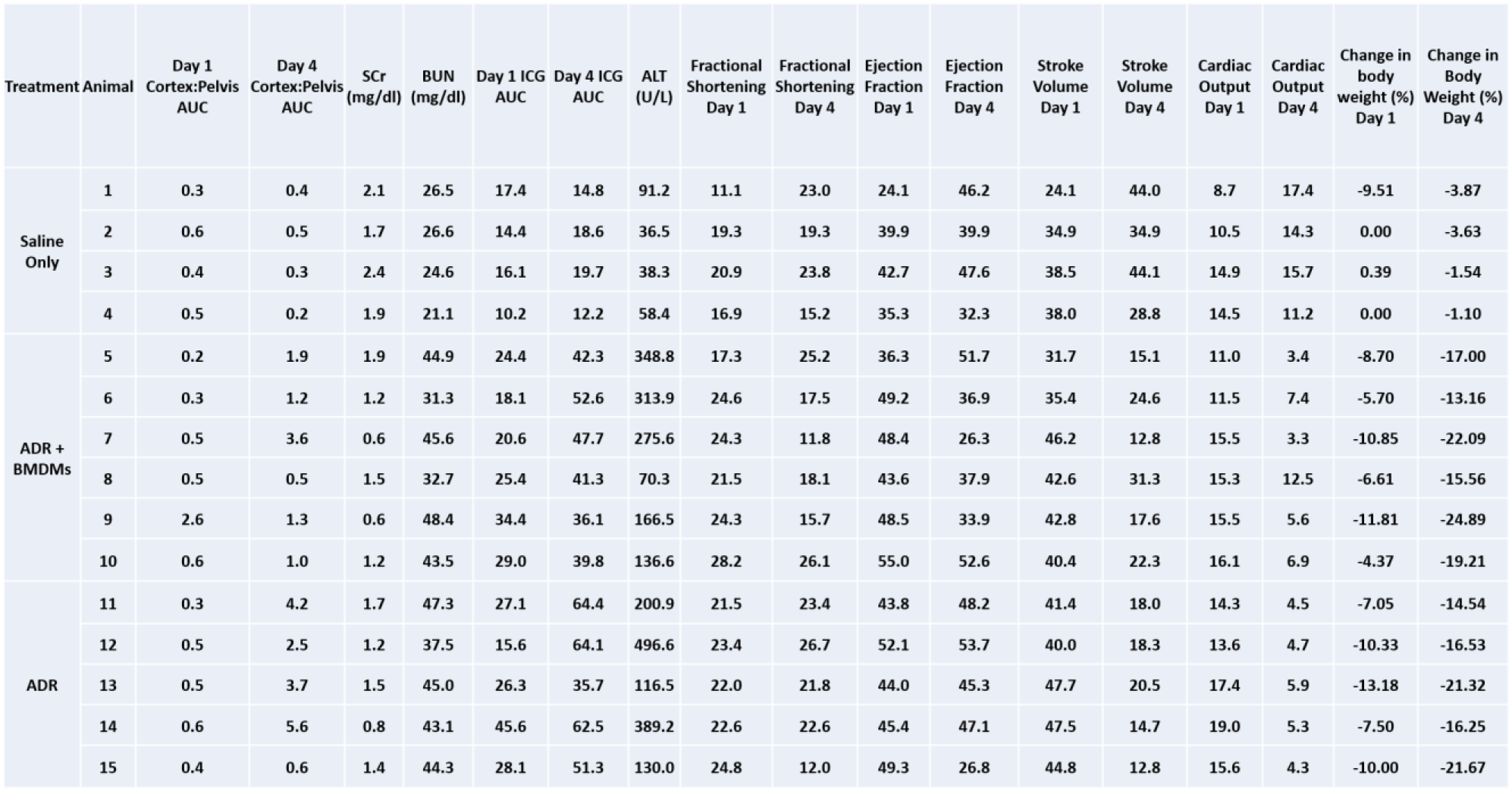
Summary of all imaging and biomarker analysis for individual mice

